# Topological dependent modifications of gene expression in the preeclamptic placenta under aspirin treatment

**DOI:** 10.64898/2026.10.23.761468

**Authors:** Ikram benouda, Daniel Vaiman, Francisco Miralles, Paul Guerby, Géraldine Gascoin, Charlotte Dubusc, Josépha Attal, Sofia Gonzalez Matatoros, Céline Méhats, Stefan R. Hansson

## Abstract

Preeclampsia is a hypertensive disorder of pregnancy characterized by systemic endothelial dysfunction and placental inflammatory activation. Low-dose aspirin reduces preeclampsia incidence when initiated early, yet its molecular effects on the diseased human placenta remain poorly defined. We performed a topologically resolved transcriptomic analysis of placentas from two independent cohorts of preeclamptic women treated or not with aspirin. Placental samples were obtained either from the villous surface (syncytiotrophoblast-rich) or from deep parenchymal regions (stromal and endothelial-rich).

In villous samples, microarray profiling identified 1,836 differentially expressed genes, predominantly down-regulated. Gene-set enrichment demonstrated marked suppression of pathways central to hypertensive placental pathology, including TNFα/NF-κB signaling, hypoxia-responsive programs, glycolysis, and estrogen-response cascades. RNA-seq analysis of parenchymal samples identified 645 differentially expressed genes, again dominated by down-regulation of cell-cycle progression, mTORC1 signaling, inflammatory pathways, and metabolic stress responses. Across cohorts, 116 genes were consistently down-regulated by aspirin and enriched for hypoxia, NF-κB, estrogen-response, and inflammatory signatures. Promoter analysis implicated transcription factors central to vascular inflammation and stress signaling, including NF-κB1, RELA, HIF1A, PPARG, and PARP1.

Functional validation in BeWo trophoblasts confirmed aspirin-dependent modulation of key placental transcripts and altered secretion of MMP9, ANGPT2, and soluble endoglin (sENG) molecules directly implicated in endothelial dysfunction and hypertensive disease.

These findings demonstrate that aspirin attenuates core inflammatory, hypoxic, and metabolic pathways across multiple placental lineages in preeclampsia, providing a mechanistic framework linking early aspirin administration to improved vascular and placental adaptation in hypertensive pregnancy. The action of aspirin upon differentiation of trophoblasts contribute to explain why aspirin protects only when administered early in pregnancy, before the end of the first trimester.

## Introduction

Preeclampsia is a common pregnancy disorder characterized by de novo gestational hypertension associated with proteinuria, according to the classical definition ^1^. It involves generalized endothelial activation and can have long-term consequences for both maternal and fetal health in humans ^2^, as well as in animal models ^3,4^.

Aspirin (acetylsalicylic acid) is one of the oldest drugs used in human medicine. The antipyretic and anti-inflammatory properties of willow bark, which contains salicylates, were likely exploited more than 3,500 years ago ^5^. In the field of hypertensive disorders of pregnancy—and particularly preeclampsia—there is extensive scientific literature on aspirin as a therapeutic agent (1,970 PubMed entries combining “preeclampsia” and “aspirin” as of september 2026). Low-dose aspirin (typically 75– 150 mg/day) has been used for more than a decade in many countries, especially in women who experienced preeclampsia during their first pregnancy.

In the 1980s, several studies reported potential benefits of aspirin in preventing preeclampsia ^6,7^. These early investigations involved small cohorts and produced inconsistent results. Meta-analyses nevertheless indicated a modest but significant protective effect. For example, Sibai and colleagues reported a reduction in preeclampsia incidence from 6.3% to 4.6% among women receiving aspirin ^8^, although the dose used (60 mg/day) was relatively low compared to most studies where the dose was comprised between 75 and 150 mg/day, while a recent studies did not indicate obvious differences between 81 and 162 mg/day within at risk populations ^9^. Other studies failed to show benefit ^10^.

A major turning point came with the meta-analysis by Bujold et al. ^11^, which demonstrated that the timing of aspirin administration is critical. Initiation before 16 weeks of gestation markedly reduced the risk of preeclampsia (relative risk 0.47), and even more so for severe forms (relative risk 0.09). In contrast, aspirin started after 16 weeks had no significant effect. These findings established early aspirin administration as an effective preventive strategy. This conclusion was reinforced by a large randomized controlled trial published in 2017 ^12^, showing a reduction in preeclampsia incidence from 4.3% to 1.6% when aspirin was appropriately administered. The beneficial effects of early low-dose aspirin have since been confirmed in various populations, including high-risk cohorts such as the Blida population, where preventive treatment at 8–10 weeks proved effective ^13–15^.

Aspirin’s most widely recognized mechanism of action involves modulation of inflammation and anti-platelet effect. As summarized by Angiolillo et al. ^16^, aspirin irreversibly acetylates prostaglandin synthases PTGS1 (COX-1) and PTGS2 (COX-2) by acetylation at Ser530 and Ser516, respectively ^17^. PTGS1 is inhibited at low doses, whereas PTGS2 requires higher doses. These enzymes generate prostaglandin G2/H2, the precursor of thromboxane A2 (a vasoconstrictor) and prostacyclin (a vasodilator), leading to complex and sometimes opposing physiological effects. Acetylated PTGS2 also promotes the formation of aspirin-triggered lipoxins and resolvins, which actively contribute to the resolution of inflammation ^18,19^.

Beyond PTGS inhibition, aspirin exerts additional effects. Its metabolite salicylate inhibits IKKβ, a kinase required for NF-κB activation ^20^, and can activate AMPK, a key regulator of fatty acid oxidation, glucose uptake, and mitochondrial biogenesis ^21^. Thus, aspirin influences multiple cellular pathways.

Despite these advances, many questions remain unanswered, regarding aspirin’s mechanisms in the preeclamptic placenta. It is unclear whether its effects primarily target endothelial cells (placental or systemic) or whether other placental cell types—such as Hofbauer cells, trophoblasts, or syncytiotrophoblasts—are also affected. Moreover, at the low doses used for preeclampsia prevention, the specific metabolic pathways modulated by aspirin remain uncertain. The lack of efficacy when aspirin is administered after 16 weeks is particularly intriguing: if its main actions were anticoagulant or anti-inflammatory, one would expect benefits even later in gestation or after the onset of the symptoms. This suggests that aspirin may act on early trophoblast populations, possibly modulating trophoblast properties (invasion, proliferation), topics that have been insufficiently studied. Given that the placenta is central to the pathophysiology of preeclampsia, we focus here on this organ.

Using transcriptomic analyses, we characterize the effects of aspirin on the preeclamptic placenta in two independent cohorts of women treated or not with aspirin, with samples collected from distinct placental regions. In the first cohort (from Lund, Sweden, analyzed with Microarrays), the samples were taken from the maternal side of the placenta (facing the uterus), and correspond mainly to villous trees lined by the syncytiotrophoblast. In the other cohort (from Toulouse, France, analyzed by RNA sequencing), the samples were from carrots taken inside the placenta.

We further integrate recently published single-cell RNA-seq datasets through deconvolution analyses to infer how aspirin may act on specific placental cell subtypes. Finally, we validate part of these findings in the BeWo trophoblast model at both transcriptomic and secretomic levels. To our knowledge, this is the first systematic study of aspirin-associated gene expression changes in placentas from preeclamptic women.

## Material and Methods

### Human samples

Samples were collected at delivery after the patients gave their informed consent. The clinical characteristics available are presented as **Table 1**.

**Table 1:**
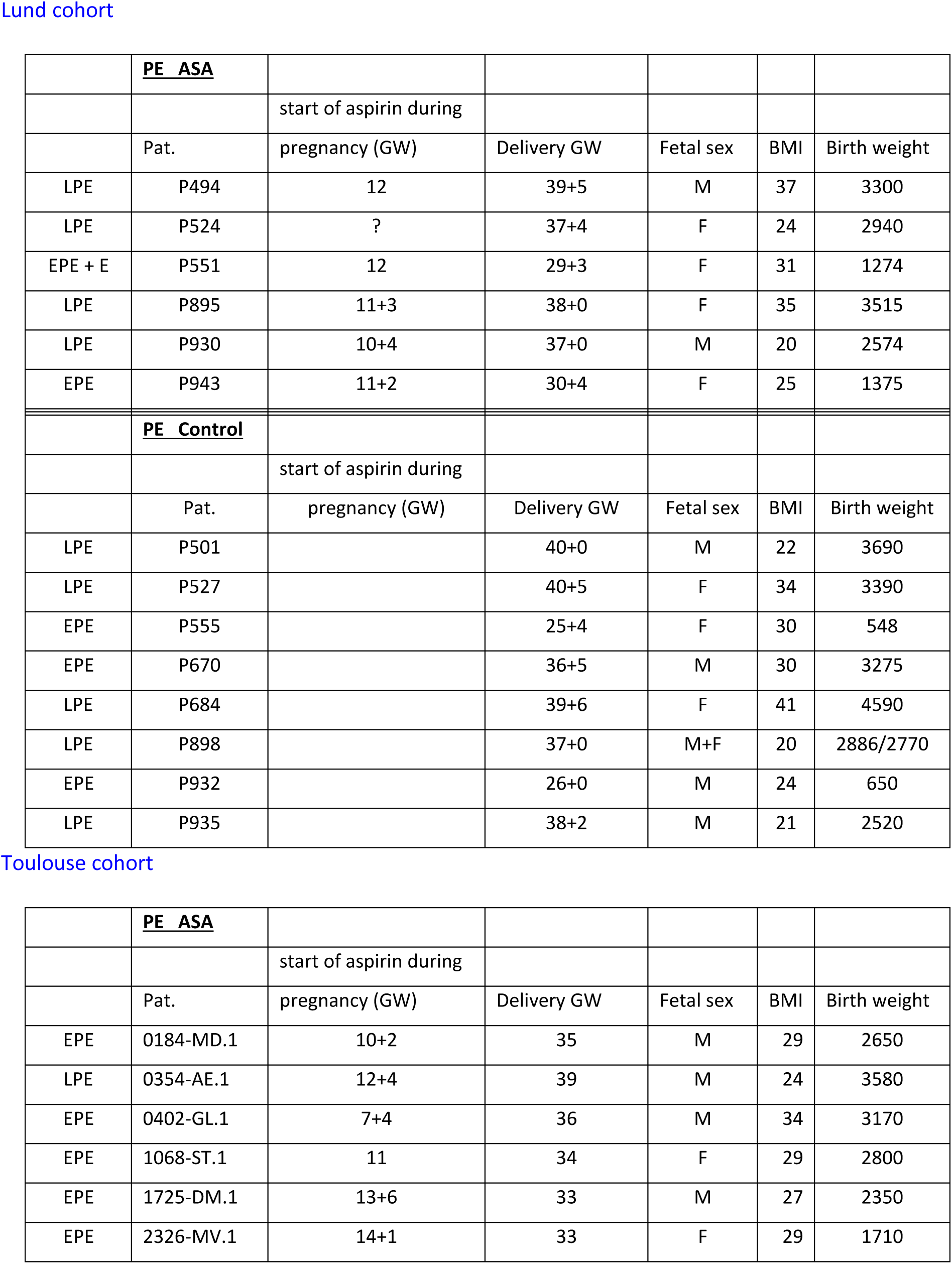
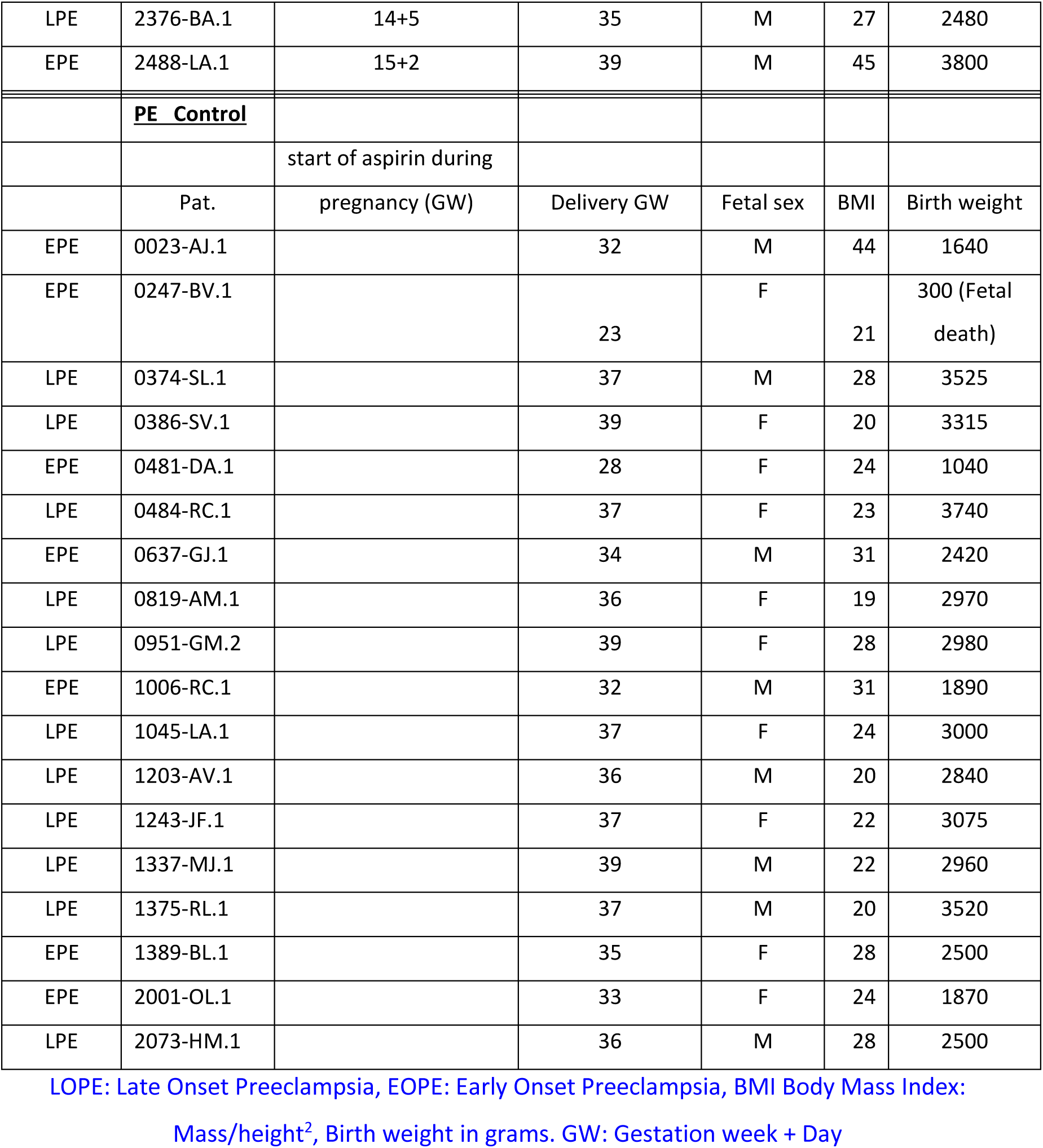
Patient characteristics.

The patients of the first cohort (all are preeclampsias, 6 from patients treated with aspirin and 8 from untreated patients) were collected according to the recommendations of the local Ethical committee at Lund University, LU 803-2. Three 1 cm3 samples were collected from the maternal side of the central placenta after removal of the fetal membranes and washing the samples extensively in PBS. The samples were then snap-frozen in liquid nitrogen and kept at -80°C before processing. The Toulouse cohort was composed of 1cm3 samples collected from the inside of the placental tissue (fetal face). They included placental samples from 26 preeclamptic patients, 8 of which were treated by aspirin, and therefore 18 without treatment. This samples are part of a specific collection from the ‘Centre de Recherches Biologiques’ inside the Institut Universitaire du Cancer de Toulouse, and were collected from patients following signed informed consent and local regulations (iuct-oncopole.fr). The histological origin of the samples is presented in Figure 1A. The cohorts taken as a whole were thus composed of placental samples from 40 preeclamptic women, 14 treated with aspirin and 26 untreated.

**Figure 1:**
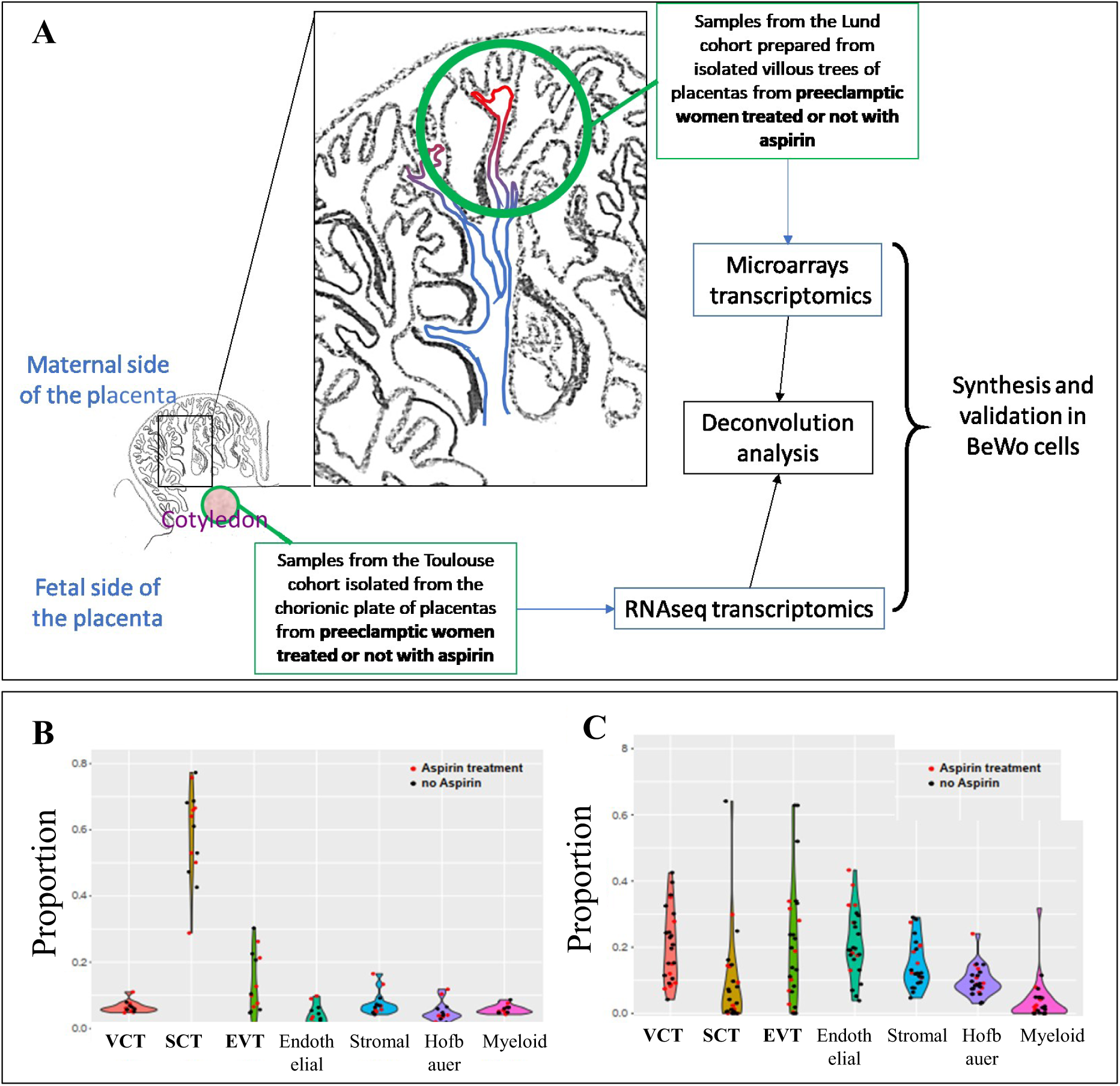
Protocol of the study and deconvolution analysis. **(A)** Placental samples were obtained from villous structures dissected and throughly washed (Lund cohort), or from the inside of placental cotyledons (Toulouse cohort). Transcriptomics was performed, deconvolution validated the origin of the samples and deregulated genes following aspirin treatment were identified and some validated in the BeWo trophoblast model. The transcriptomic data were submitted to deconvolution analysis using the CibersortX algorithm. This allowed to identify the proportions of the main placental cell-types in the samples of first **(B)** and **(C)** second cohort. The red dots represent the samples issued from preeclamptic pregnancies who received aspirin, while the black dots correspond to the placental samples from preeclamptic women without aspirin administration. HB stands for Hofbauer cells, SCT for syncytiotrophoblast, EVT for extravillous trophoblast and VCT for villous trophoblast^23^.

### Microarray analysis (first cohort, Lund)

After validation of the RNA quality using a 2100 Bioanalyzer analysis (Agilent), all the samples were shown to have a RNA Integrity Number (RIN) >7, and could then be used for microarray analysis. One hundred ng of RNA from each placental sample were analyzed by ClariomD (Affymetrix) microarray assay, at the Genom’IC platform of the Cochin Institute. This tool interrogates 133,847 probes, including coding genes (18,858), non-coding genes (66,845) and genes deduced by bioinformatics analysis (10,001), in addition to pseudogenes, small RNA and ribosomal RNAs. Overall, more than 540,000 transcripts were analyzed considering splicing variants. (.CEL) files were processed using the Transcriptome Analysis Console (TAC) software (Thermo Fisher Scientific), performing the default Robust Multi-Array normalization (RMA). This algorithm performs background adjustment, followed by quantile normalization and summarization. The results entitled "Placental Effects of Aspirin treatment in preeclamptic women" have been assigned the ArrayExpress accession number E-MTAB-12434 in the EBI-EMBL repository.

### RNA-seq analysis (second cohort, Toulouse)

The RNA seq was performed at BMKGENE company from RNA sent after quality validation. In a folder presented as a Supplementary document are available the following documents (**Supplementary dataset 1**). The available folders are shown as **Table 2**

**Table 2:**
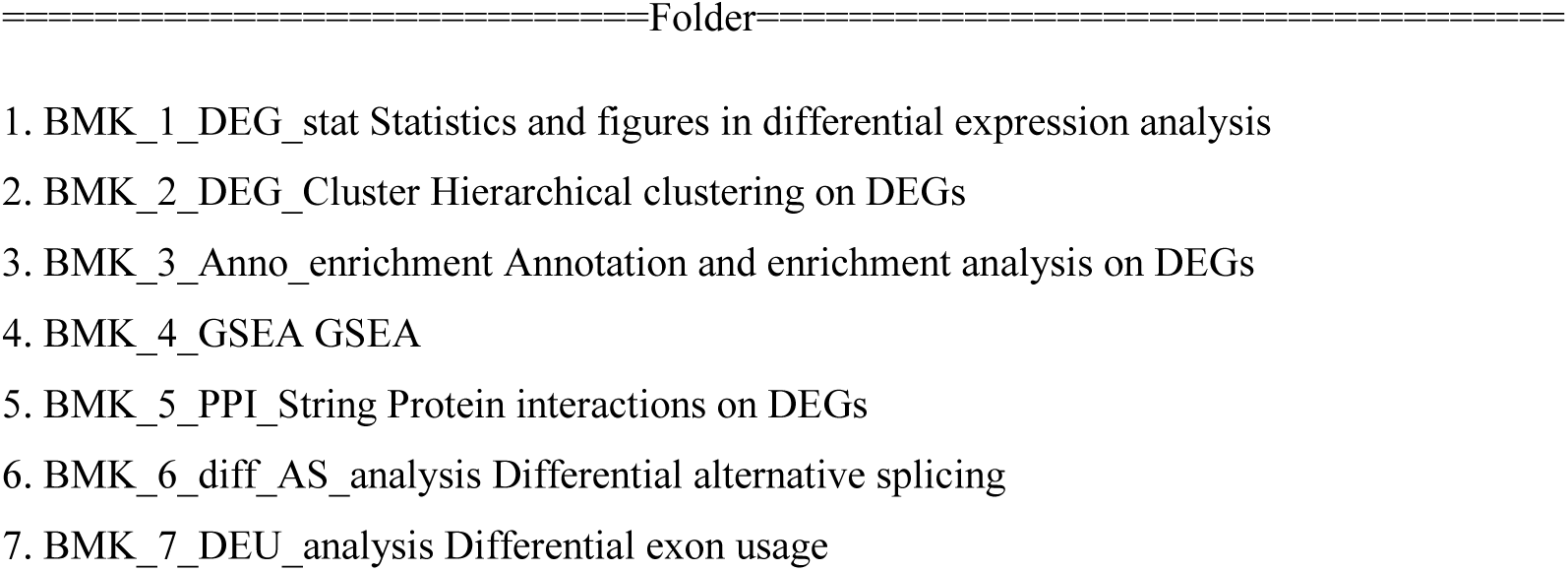
Folders provided describing the RNA seq data provided by BMKGENE.

### Deconvolution

Computational deconvolution methods allow to extract cell type-specific information from gene expression data obtained from heterogeneous tissue samples. We submitted our bulk transcriptomic data from both cohorts, to the deconvolution algorithm CibersortX to characterize the cell types, and their proportion ^22^. CibersortX uses a reference single-cell RNA-seq experiment to deconvolute bulk microarray data. The reference used in our study consisted of a single-cell dataset experiment conducted on the human placenta ^23^.

### GSEA analysis using Webgestalt

The list of genes, associated with their relative induction levels (preeclampsia versus preeclampsia with aspirin) were pasted into the Gene List window of WebGestalt (www.webgestalt.org). The organism of interest was Homo sapiens, the Method used GSEA, and the databases screened was mainly \community-contributed\Hallmark50’.

### BeWo cell culture and treatment

BeWo cells were cultivated in F12 medium (Life Technologies) supplemented with 10% fetal bovine serum (FBS) and 1% penicillin/streptomycin in 6-cm diameter plates, up to 60% confluence; Cells from two different experiments were cultivated in 12-well plates. Effect of aspirin was tested at several doses and 18 mg/L was found optimal in terms of response without toxicity based upon our data and the literature, corresponding to ∼0.1 mM. For the forskolin-treated samples, forskolin (20μM final) was added on day2, and the RNA was collected on day 5 in TriZol, followed by standard protocols for RNA extraction.

### RT-qPCR and primer choice

All the qPCR primers used in this study were defined from the Harvard collection of qPCR primers, using the website https://pga.mhgh.haravrd.edu/primerbank/. qPCR was performed in 15 μl using SyberGreen 2X kit from BiolIne, using a LightCycler480. Each qPCR reaction was followed by a melting curve analysis to check for the homogeneity of the PCR products. Normalization was done using PPIB (peptidylprolyl isomerase B, cyclophilline) as a reporter gene, after having checked the excellent stability of the gene expression in our cell cuture model.

### Luminex analysis

Cell supernatants were analyzed with the Bioplex200 device from Bio-Rad with Human Magnetic Luminex Assays Plex from R&D Systems (Bio-Techne, Minneapolis, MN) according to the manufacturer’s instructions. Standard curves were calculated from the standard samples, and the values fitted to a Logistic-5Pl curve; this allowed us to estimate the concentrations of the relevant samples

## Results

### 1. Cell composition analysis of samples from the two cohorts, assessed by deconvolution, revealed that each cohort present different proportion of cell types, in accordance with the sampling zone

To analyse the coherence of the cell composition with the site were the sample was taken from in the placenta and that differed between the two cohorts, we performed a deconvolution analysis to evaluate sample variation inside each dataset. To this end we used as reference the placenta scRNA-Seq data ^23^ and the deconvolution algorithm CibersortX ^22^, as presented in the Methods section.

The deconvolution analysis of the first cohort (Figure 1B) showed that these samples contained on average a very large proportion of SCT (58%), and a high proportion of EVT (13%). The content in VCT was small (6%). Immune cells were present as well as Myeloid cells (5.7%) and Hofbauer cells (5%). Finally, the samples contained also stromal cells (7%) and a small proportion of endothelial vascular cells (3.5%). This is consistent with the known cell composition of the villous tree.

The deconvolution analysis of the second cohort (Figure 1C) showed that the samples contained a very small mean proportion of SCT (9.3%), and a higher proportion of EVT (20%), and VCT (19%). The samples were also enriched in fibroblasts (15%) and endothelial vascular cells (21%). The contribution of immune cells was from Myeloid (3.7%) and Hofbauer cells (9.6%), in accordance with the origin of the placental sample taken inside the mass of the tissue.

In both cohorts, no significant differences in cell composition were detected between the samples of aspirin-treated or untreated women. Therefore, the deconvolution analysis indicates that the cell composition of the second cohort differs significantly from the first one. Consistently with the sampling procedure, since the first cohort was obtained from washed chorionic villi, mostly at the villous surface (floating villi) rather than deep basal plate/anchoring regions. On the contrary the proportions observed int the second cohort are consistent with the sampling carried out from villous parenchyma of the placental disc, enriched in anchoring villi at the maternal–fetal interface ^24^. This massive discrepancy was exploited to separately analyse the transcriptomic effects of aspirin on two different topologic regions of the human placenta.

### 2. Comparison between aspirin-treated versus non-treated women in the first cohort: transcriptome of the chorionic villi

#### a. Global bulk analysis

At a threshold of p < 0.05, and excluding the sex chromosomes, 5429 tags were identified in the Affymetrix Clariom microarrays, that were differentially expressed following aspirin treatment, and 1836 were referred to as genes (excluding the genes identified by bioinformatics in the ‘Aceview’ category of the array); 635 genes were up-regulated (34.7 %), and 1201 (65.3%) were down-regulated, consistently with the excess of down-regulated transcript in the placenta under aspirin treatment that we described in mice ^25^. These genes were distributed along categories described in **Table 3**.

**Table 3:** Deregulated transcripts from the ClariomD microarray.

| Group | Total | Passed Filter | Up-Regulated | Down-Regulated |
| --- | --- | --- | --- | --- |
| Multiple_Complex | 29510 | 876 | 223 | 653 |
| NonCoding | 66845 | 576 | 276 | 300 |
| Coding | 18858 | 166 | 28 | 138 |
| Pseudogene | 4340 | 67 | 34 | 33 |
| Small_RNA | 2386 | 67 | 20 | 47 |
| Precursor_microRNA | 3297 | 52 | 34 | 18 |
| Unassigned | 10001 | 17 | 10 | 7 |
| Ribosomal | 507 | 15 | 10 | 5 |
| tRNA | 6 | 0 | 0 | 0 |

**The data were systematically** analyzed by a Gene-Set Enrichment Analysis approach (GSEA ^26^), using Webgestalt (https://www.webgestalt.org/) ^27^. At a first level of analysis, we performed this analysis against the 50 ‘Hallmarks’ defined previously by the Broad Institute (https://www.gsea-msigdb.org/gsea/index.jsp). These hallmarks summarize, with minimal redundancy, 50 clusters of genes recapitulating important cellular and biological functions (accessible at the following Broad Website http://www.gsea-msigdb.org/gsea/msigdb/genesets.jsp?collection=H). In **Figure 2A** are summarized the major results of this analysis. Thirty thousand genes were unambiguously identified from the microarray probes and interpreted by WebGestalt. Consistently with the overrepresentation of down-regulated genes in the array, no significant cluster of up-regulated genes could be found with a False Discovery Rate (FDR) set at <= 0.05. Several strikingly enriched cascades where directly connected to inflammation, such as ‘TNFα signaling via NFκB’, where 95 genes out of 191 in this cascade contribute to the density of down-regulated genes. These genes with their level of down-regulation are presented, as well as other leading-edge gene list for enriched hallmarks 2 to 6 in **Supplementary Table S1.**

**Figure 2:**
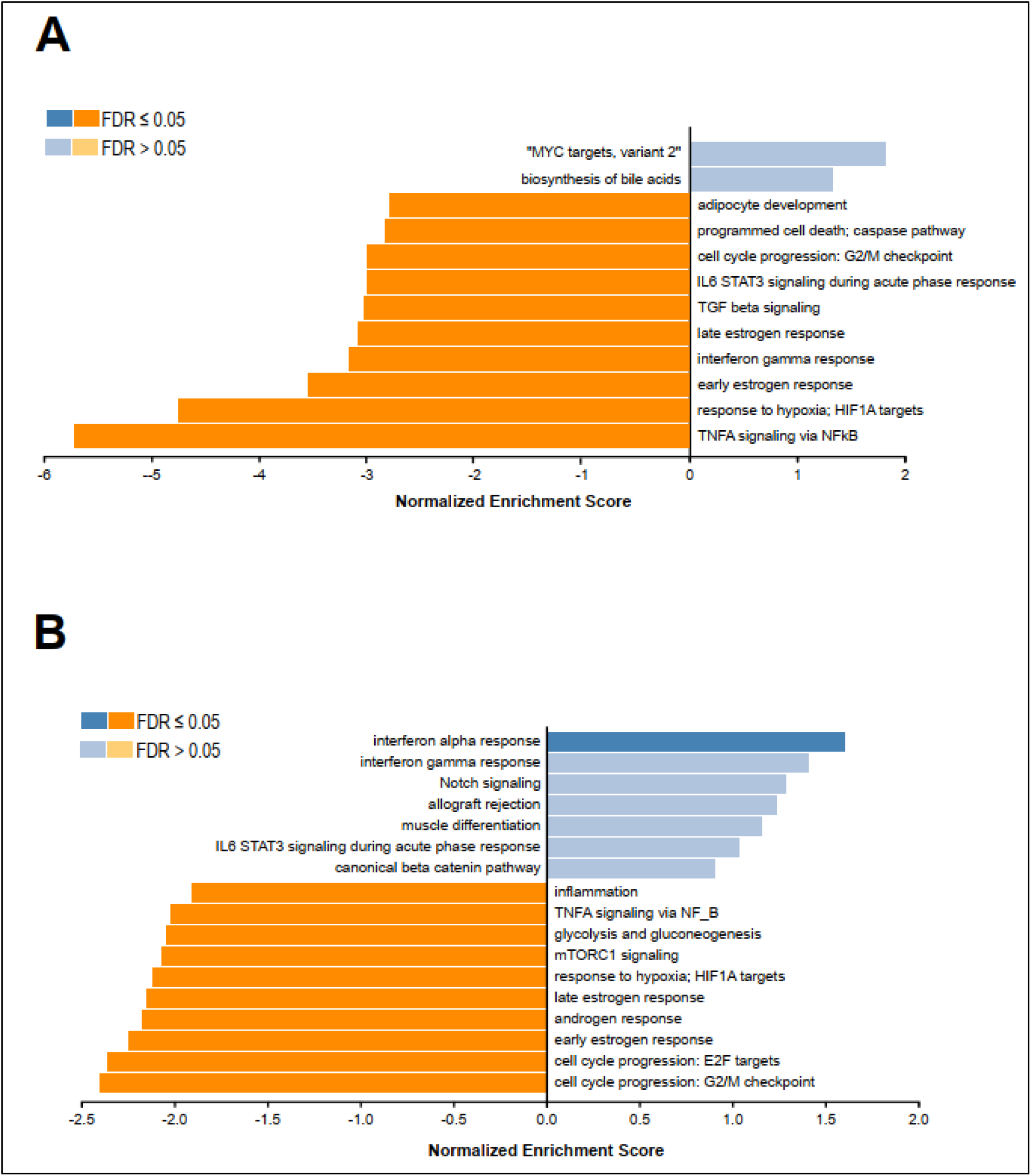
**Gene Set Enrichement Analysis (GSEA)** using the molecular signatures database Hallmark50 gene set collection (www.gsea-msigdb.org/gsea/msigdb/collections.jsp). (A) GSEA of the first cohort. (B) GSEA of the second cohort. The significant hallmarks (FDR ≤0.05) are shown in dark orange.

#### b. Cell-specific analysis of the transcriptomic effects of aspirin on the first cohort

To determine whether aspirin treatment affected gene expression in specific placenta cell types we re-analyzed by GSEA the different cell types separately (Figure 3A).

**Figure 3:**
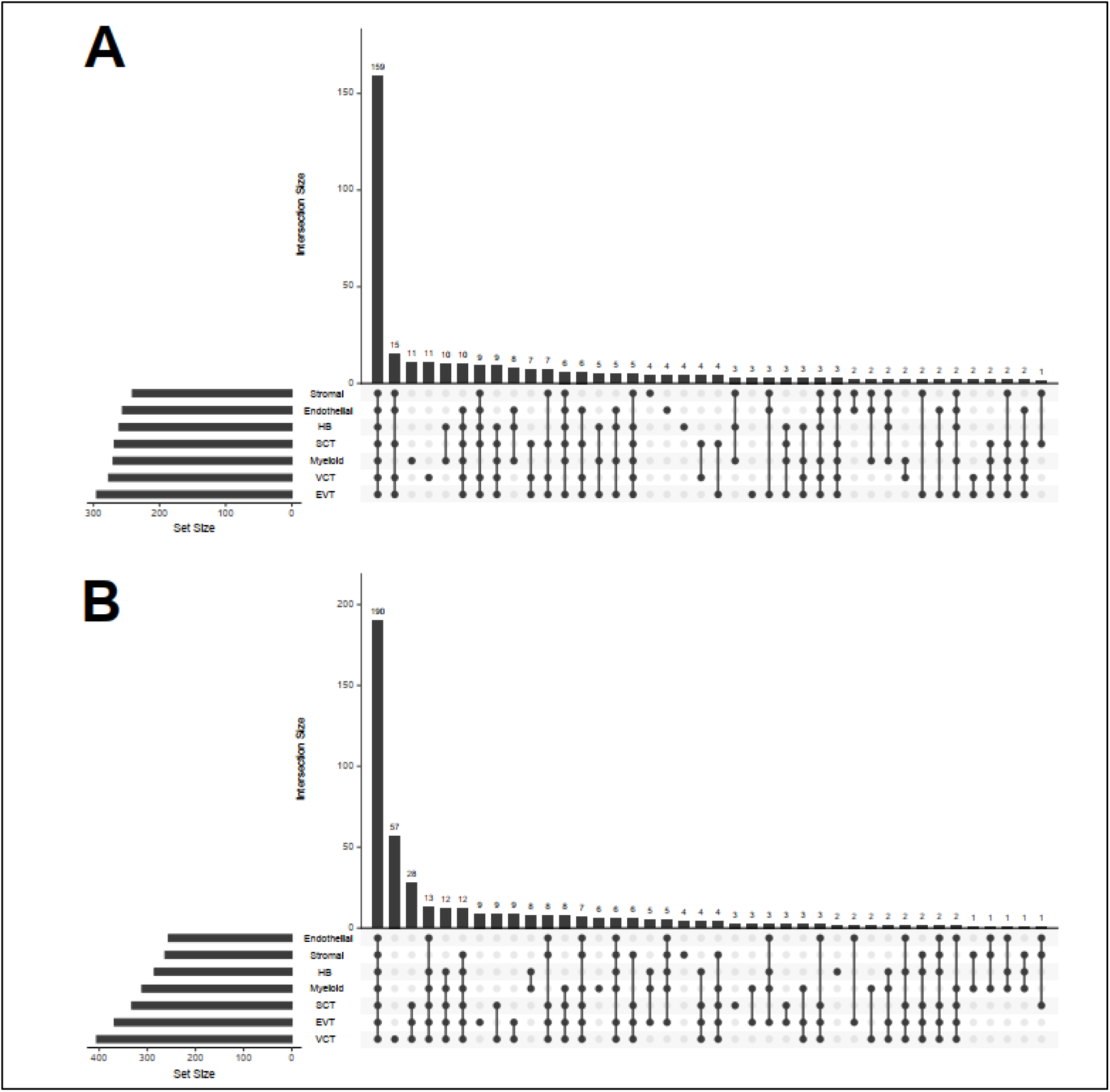
**UpsetR plots summarizing the predicted placental expression** of the down-regulated genes associated with the enriched hallmarks identified by GSEA on the first cohort (A) and second cohort (B). Cell-type placental expression was obtained from a published single-cell RNA sequencing data set on human placenta at term. HB stands for Hofbauer cells, SCT for syncytiotrophoblast, EVT for extravillous trophoblast and VCT for villous trophoblast ^23^.

The genes in the leading edge of the enriched Hallmarks were confronted with the placenta cell type signatures to infer their spatial expression This analysis indicates that many of the aspirin down-regulated genes are shared in common between the different cell types of the placenta. This includes genes such as: JUN, ATF3, ALDOA, LDHA, GPX1, PHLDA2, RELA, SERPINE1 and NFKB1. Therefore, many aspirin down-regulated genes are expressed in more than one cell type, indicating that aspirin treatment does not affect a single cell lineage. To further explore the effects of aspirin on the placenta we took advantage of the scRNA-Seq analysis comparing last trimester severe preeclamptic placentas and matched non-pathological placentas ^23^. We used this data to generate gene sets of modified gene expression in each cell-type of the preeclamptic placenta. Then we performed GSEA on our expression data against these preeclampsia cell-type specific gene sets. Very significant enrichments (FDR< 0.001) were detected in the negative phenotype, that is against the gene sets representing genes up-regulated in preeclampsia who were down-regulated in our microarray following aspirin administration (**Supplementary Figure 1**). The highest Normalized Enrichment Scores (NES) were obtained by the gene sets composed of genes up-regulated by preeclampsia, and down regulated specifically by aspirin administration, in the SCT (NES= -3.01) and VCT (NES= -2.84), followed by the Hofbauer cells (NES= -2.58) and Myeloid cells (NES= -2.57). Significant enrichments were also detected with the Endothelial (NES= -2.55), Stromal (NES= -2.37) and EVT (NES= -2.08) gene sets. This analysis confirms that aspirin administration down-regulates specifically genes that are up-regulated in the placenta by preeclampsia in different cell-types of the placenta.

### 3. Comparison between aspirin-treated versus non-treated preeclamptic women in the second cohort: transcriptome of the placental mesenchyme

#### a. Global bulk analysis

The second cohort was composed of 24 preeclamptic placenta samples treated (n = 7) or not (n= 17) with aspirin, that were obtained from a carrot inside the thickness of the placenta, and thus containing a high proportion of stromal cells and much less syncytiotrophoblast. The RNA-Seq analysis of the placentas of this group showed that at a threshold of p <0.05 and absolute FC >= 1.5 a total of 645 genes were differentially expressed (DEGs). Of these 262 were up-regulated and 383 were down-regulated in the placentas of women under aspirin treatment versus the non-treated group. To investigate the functions of these DEGs we performed GSEA against the Gene Set Collection Hallmarks50 of the Molecular Signature Data Base (MSigDB) of the Broad Institute (a group of 50 gene sets defining non-redundant broad biological functions). At a FDR < 0.05, this analysis (**Figure 2B)** identified a unique up-regulated gene set: interferon alpha response (NES=1.59), and nine down-regulated gene sets. The later include gene sets related to: cell cycle progression (NES = -2.37), estrogen response (NES = -2.21), response to hypoxia (NES = -2.11), mTORC1 signaling (NES =-2.08), glycolysis and gluconeogenesis (NES = -2.08), TNFA signaling via NFkB (NES = -1.99) and Inflammation (NES= -1.90). The genes included in the leading edge of each enriched gene set and their fold changes are presented in the S**upplementary** T**able S2.**

#### b. Cell-type effects of aspirin in the second cohort

Alike what was done with the first cohort we used the cell-type specific signatures obtained from the reference scRNA-seq to infer the cell-type expression of the down-regulated genes under aspirin. As for the first cohort this analysis indicate that many down-regulated genes are expressed in common by the different cell types of the placenta (**Figure 3B**). These common genes included among others: ATF3, SERPINE1, LDHA, PLIN2, TNFAIP3, REL, TNFAIP3 and MAP3K1. However, in this cohort we observe that many down-regulated genes are expressed uniquely in the VCT. These includes genes such as: BUB1B, CDC20, CHEK1, MKI67, TOP2A. Overrepresentation analysis (ORA) showed that these genes are involved in cell cycle progression. Therefore, many aspirin down-regulated genes are expressed in more than one cell type. This corroborates the results of the first cohort indicating that aspirin treatment is not cell type specific. This cohort was also submitted to GSEA using the preeclampsia cell-type specific up-regulated gene sets inferred from the scRNA-Seq data (**Supplementary Figure 2**). Significant enrichments (FDR< 0.001) concerned genes up-regulated in different cell-types of the preeclamptic placenta and down-regulated by aspirin administration: SCT (NES= -2.19), EVT (NES= -2.16), Myeloid (NES= -2.15), Hofbauer cells (NES= -1.88) and VCT (NES= -1.65).

Thus, in both cohorts, genes normally up-regulated in the preeclamptic compared to the healthy placenta, are specifically targeted by aspirin to be down regulated.

### 4. Common transcriptional networks in the two placental compartments

**Despite the important differences in cell composition** between the cohorts we found 116 common genes of which the vast majority (101) is down-regulated and part of the leading-edge of the enriched cell-type gene sets identified by GSEA (**Supplementary Table S3**). These are genes that are up-regulated in preeclampsia and down-regulated by the aspirin treatment in both cohorts. The ORA of these 116 genes show that they are strongly associated with the biological functions that we identified previously: response to Hypoxia (FDR= 5.13 10^-27^) TNFα signalling via NF-kB (FDR= 1.01 10^-23^), Early estrogen response(FDR= 4.27 10^-9^), Late estrogen response(FDR= 5.62 10^-6^), Inflammation (FDR= 2.19 10^-5^), etc (**Figure 4A & 4B**). The ORA analysis show also that the expression of these genes is potentially regulated by the following transcription factors: NF-kB1 (FDR= 9.85 10^-10^), RELA (FDR=5.59 10^-9^), PPARG (FDR=1.55 10^-7^), PARP1 (FDR=0.0000074), HIF1A (FDR=0.0000093), etc

**Figure 4:**
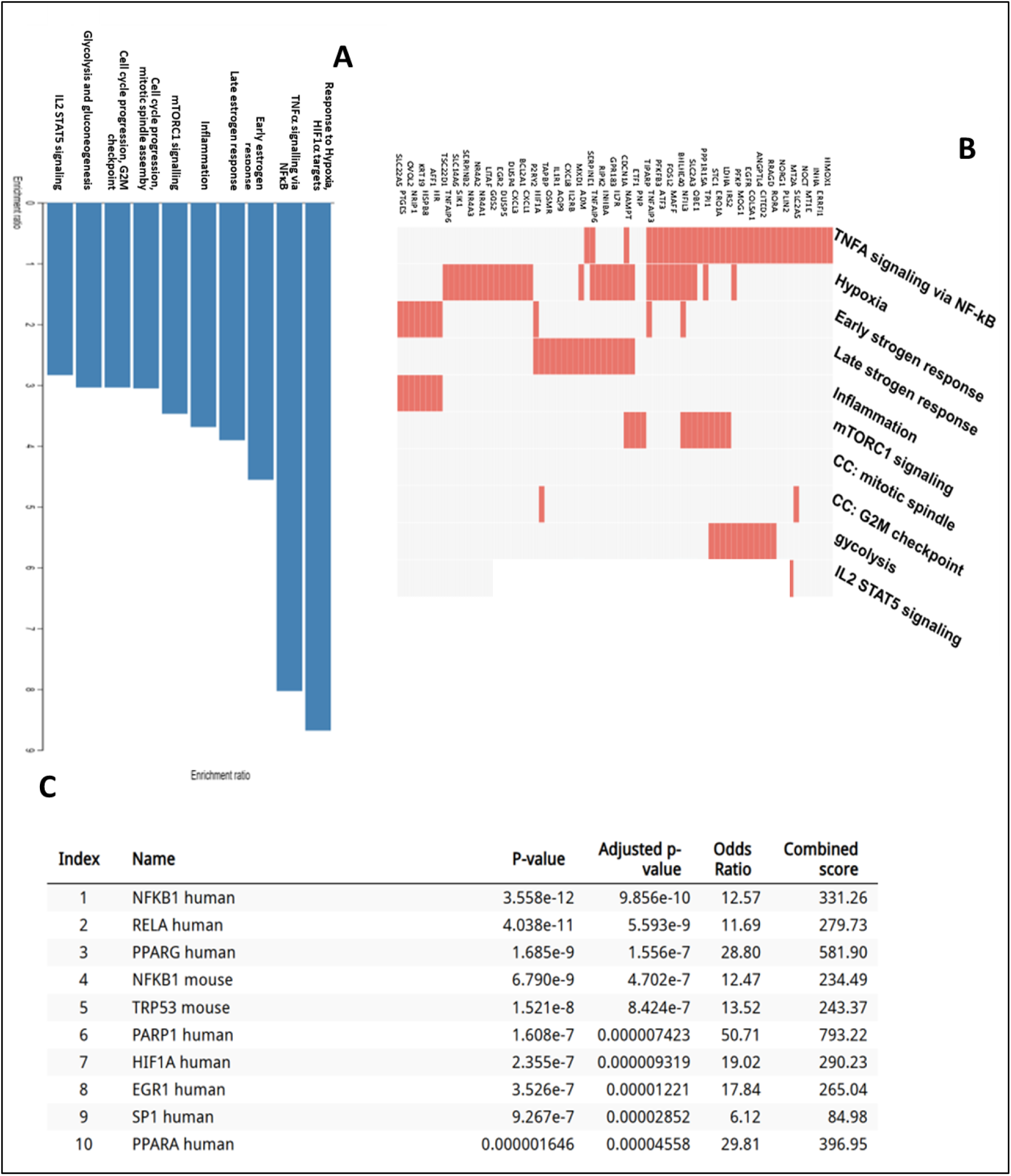
Enrichment analysis of the 101 genes which are down-regulated in both cohorts. Over Representation analysis (ORA) showing the significant hallmarks associated with the set of common down-regulated genes (A). The clustergram (B) shows a portion how these genes distribute among the enriched pathways. ORA identifying the transcription factors whose transcription factor binding sites are enriched in the promoters of the of down-regulated genes (C).

(**Figure 4C**). Also, several of the genes down-regulated in the placentas of both cohorts encode transcription factors including: ATF3, BHLHE40, NR4A1, NR4A3, CITED2, and MXD1, suggesting specific modifications in the aspirin-mediated signalling.

### 5. Mimicking aspirin effect in a fusogenic trophoblast model, the BeWo choriocarcinoma cell line

**Given the specific** aspirin effect at the placental level, we wished to check whether the trophoblast lineage that defines the placenta is itself affected directly by exposure to aspirin. For this we studied gene regulation in vitro in the BeWo model, a trophoblast cell line which is able to fuse under forskolin treatment, and thus to mimic the behaviour of villous trophoblast, as well as the behaviour of syncytiotrophoblasts. The cells were treated with aspirin and/or forskolin before proceeding to analysing RNA expression through RT-qPCR.

We analyzed in six-plicates 32 genes, 22 of which were found down-regulated by aspirin in the preeclamptic placentas (ATF3, TNFAIP3, NDRG, ADM, CRH, GATA3, INHA, TGFB1, LNPEP, ENG, LRRC1, GDF15, BAD, BCL2, PLIN1, JUN, PAPPA2, EGR1, FLT1, HPGD, MXD1, ITGA5), some that were highly relevant for studying the fusion mechanisms (SYNC1 (ERVW1) and SYNC2 (ERVFRD1), STOX1A (facilitating fusion) and STOX1B (inhibiting fusion) ^28^, as well as the regulation of apoptosis in addition to BAD and BCL2, BAX as well as proliferation KI67), and another endoretrovirus ERV3. To note, ENG and FLT1 encoding two of the principal early circulating markers of preeclampsia are included in the list.

Overall, amongst these 32 genes, 9 were expressed at a higher level following FSK treatment (ATF3, TNFAIP3, NDRG1, ADM, LRRC1, FLT1, HPGD, MXD1, ERVFRD1), seven saw there expression levels reduced (STOX1, BAX, Ki67, KRT19, PLIN1, PAPPA2 and HTRA1), and the others were either stable, or revealed an opposite effect of aspirin following the induction of fusion by FSK, especially GDF15, BAD, BCL2, JUN, ERVW1, ITGA5, and notably ENG).

In these experiments, SYNC1, SYNC2, LRRC1, GDF15, STOX1A and BCL2 were significantly induced as shown in **Figure 5**. The effect of aspirin was visible on the expression of GATA3, INHA, TGFB, LRRC1, GDF15, STOX1A and STOX1B, BAX and KI67. There was an interaction effect between aspirin and forskolin on the CRH, ENG and GDF15 gene expression. Overall, these results confirms that BeWo cells could be used to partially mimic placental effects of aspirin, as previously hypothesized, as well as for the specific response to aspirin in subsets of placental cells ^29,30^.

**Figure 5:**
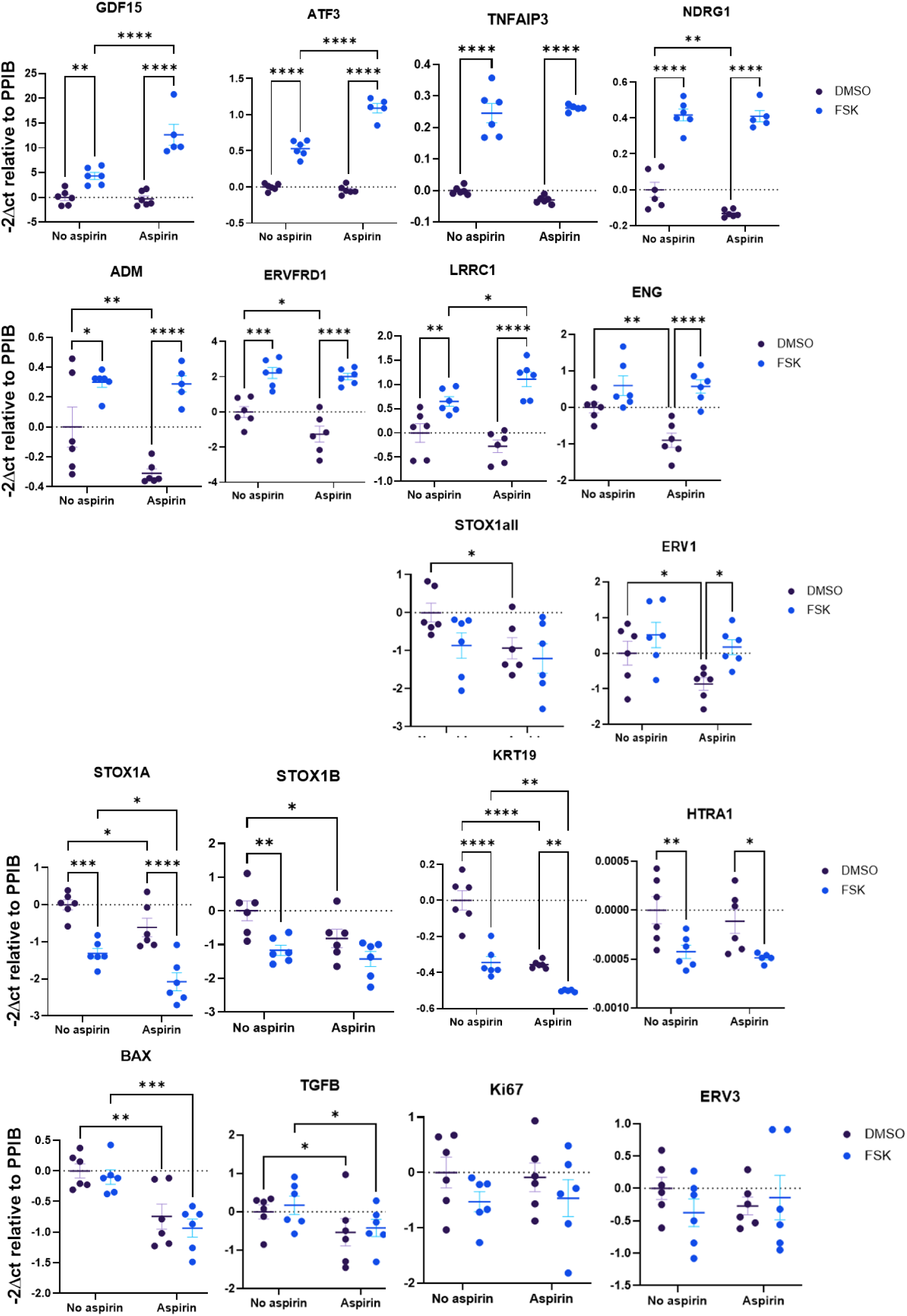
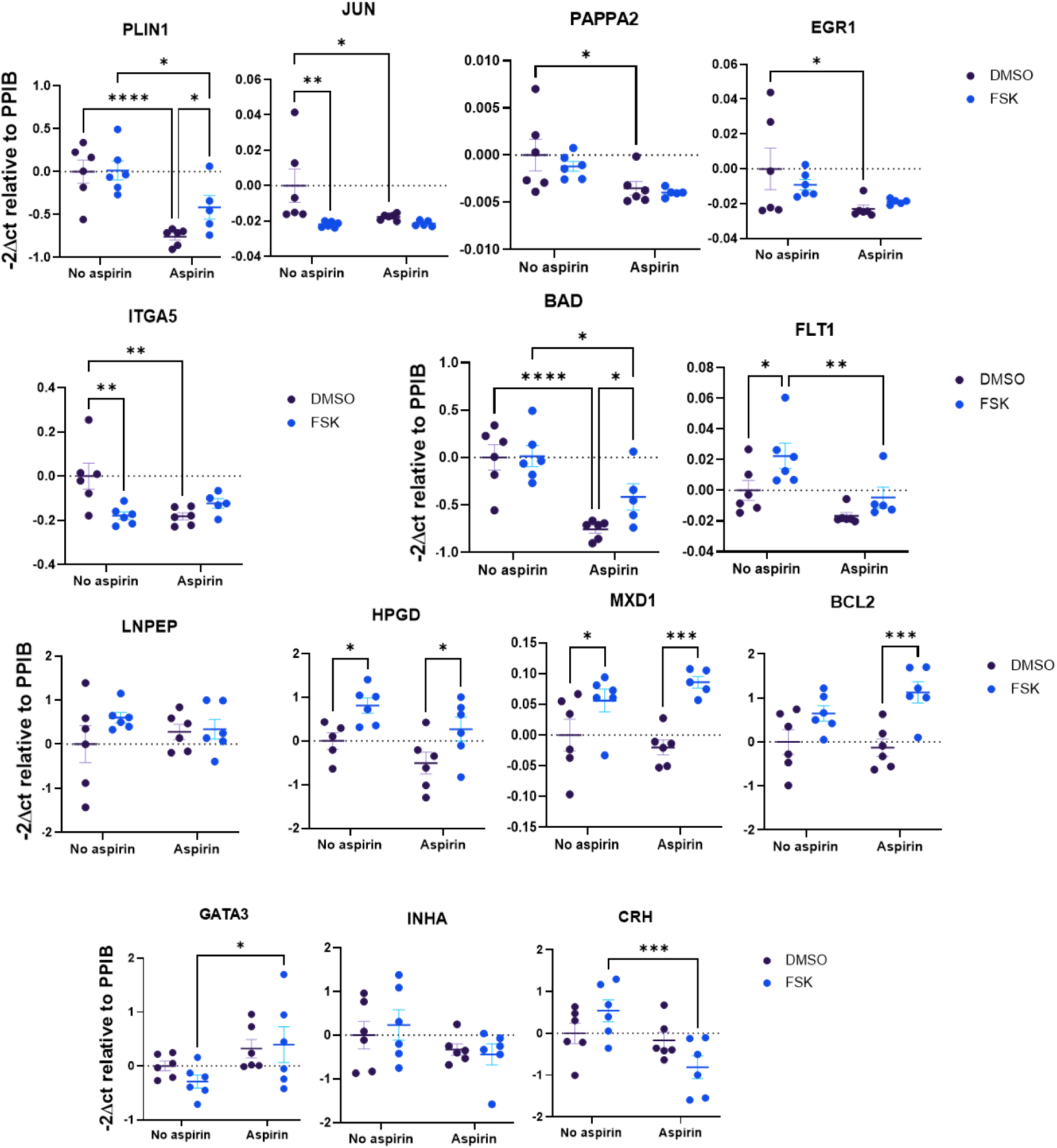
**RT-qPCR analysis of various genes identified in the placental transcriptome in the BeWo cytotrophoblast cell model**, where fusion has been induced or not with forskolin (FSK+ or no FSK), and with or without aspirin treatment (Aspi+ or No Aspi). Results are from three independent experiment carried out in 6-plicates. After significant ANOVA two-factor analysis post-hoc tests were carried out. Asterisks mark significant effects of aspirin (*: p < 0.05, **: p < 0.01, ***: p < 0.001).

This dataset of RT-qPCR confirms that FSK induces syncytialisation. Two-ways ANOVA revealed the significant induction of classical trophoblast fusion markers, SYNCYTIN1 and SYNCYTIN2, ENG, or GDF15, data that validate the model. Besides, we observe that in cells not treated with FSK, aspirin alone already pushes several trophoblast genes toward a differentiated phenotype. This is the case for GATA3, CRH, SYNCYTIN1, SYNCYTIN2, TGFB, ENG, GDF15, STOX1A and STOX1B. This suggests aspirin is not acting solely through its classical anti-platelet mechanism but directly modifies trophoblast transcriptional programs. Such observations are consistent with the absence of efficiency of late aspirin treatment in preeclampsia. CRH and GDF15 are strongly induced by aspirin treatment in DMSO BeWo cells, both able to act as enhancers of trophoblast differentiation and endocrine maturation ^31,32^. About the connections between FSK-induced fusion and aspirin treatment, the data indicates additive effects in most markers, such as INHA, GATA3, SYNC1 and SYNC2, as well as TGFB. Some presented a significant p-value for interaction, CRH, ENG, LRRC1 and GDF15. The analysis of gene expression carried out by Leavay and coworkers ^33^ showed that LRRC1 is induced in the preeclamptic placenta, as well in the seminal Nishizawa article ^34^.

**To go beyond the transcriptional** deregulation, we attempted to evaluate differentially secreted molecules in BeWo cell cultures using a Luminex panel. Amongst the protein that were detectable in the supernatant, seven had a detectable level, and are presented as **Figure 6**, in a two factors ANOVA. As shown, the fusion effect mediated by Forskolin was significant for leptin (LEP) and inhibin. A significant effect of aspirin was found on MMP9; an interaction effect was detected for Angiopoietin 2 and for Endoglin (ANGPT2 and ENG).

**Figure 6:**
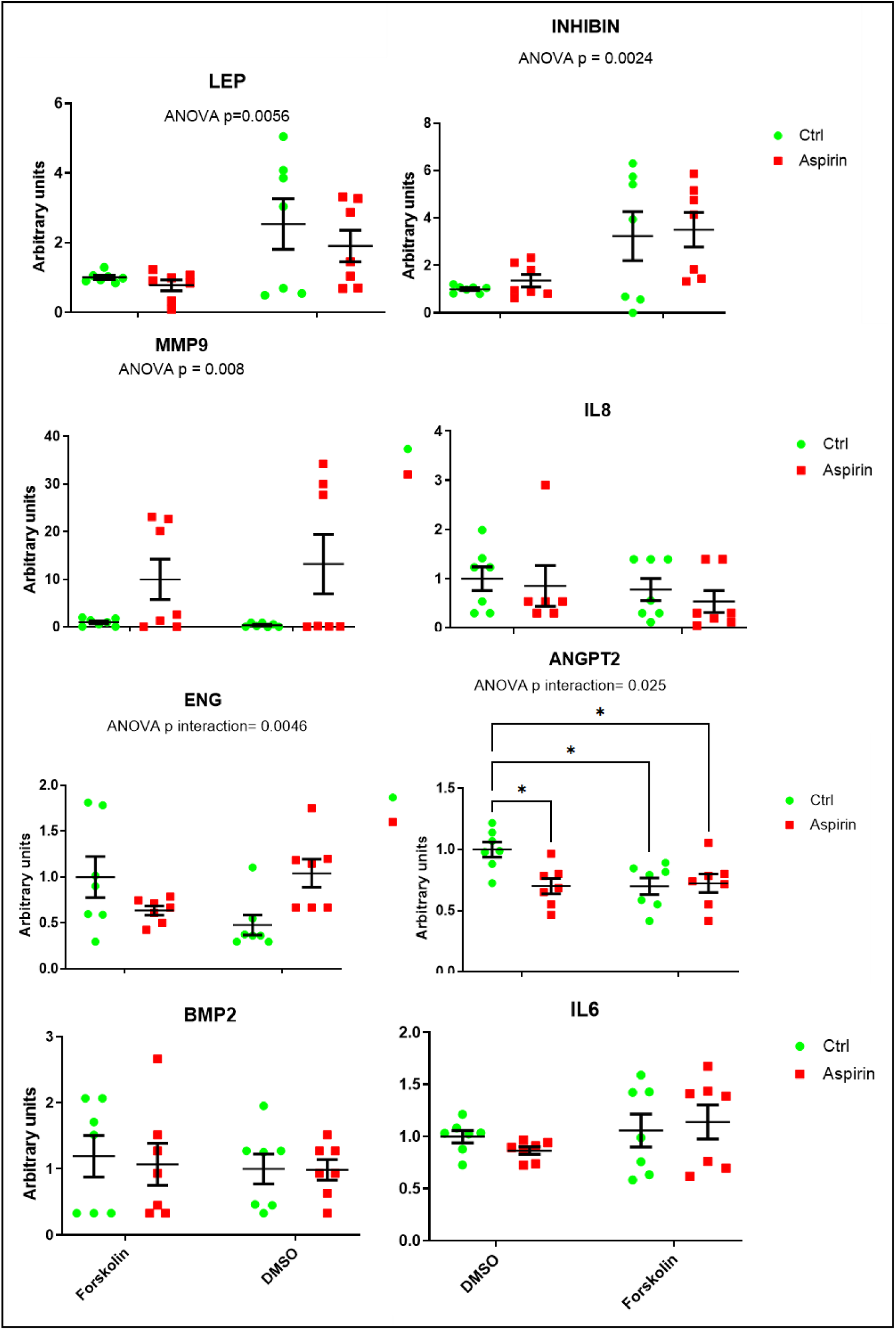
**Luminex analysis of the secretome of BeWo cells, for seven relevant proteins**, IL6/IL8 (markers of inflammation), LEP (leptin, marker of preeclampsia in the placenta), ANGPT2 (deregulated in preeclampsia ^61^ Inhibin (marker of fusion) and soluble Endoglin, ENG (a risk factor for preeclampsia), MMP9 marker of invasion and BMP2, a marker of the TGFb family reported altered in early onset preeclampsia ^62^. Results are from two independent experiments, in technical triplicates.

## Discussion

This study is, to the best of our knowledge the first to compare systematically the transcriptomics effects of aspirin on the human preeclamptic placenta. Our transcriptomic analysis was conducted on two independent cohorts of pregnant women treated with a low dose of aspirin (from 8-12 weeks of gestation till delivery). The two collections of samples were obtained from different parts of the placenta (the villous tree, Lund cohort) and the internal part of the placental tissue (Toulouse cohort), which allows to address separately the response of these compartments to aspirin treatment.

The study reveals a set of convergent and divergent results among cohorts. In both cohorts the majority of DEGs are down-regulated, and in both cases only the down-regulated genes are significantly associated with signalling pathways (biological hallmarks), showing that overall, **aspirin has a negative effect on gene expression**. Many of the down-regulated signalling pathways are common to both cohorts but a few are not. Also, differentially expressed genes (DEGs) only partially overlap, with about one hundred genes in common. Inference of spatial expression of the down-regulated DEGs using a scRNA-seq reference shows that the majority of the modified genes are expressed across all cell types in placenta, although their expression levels can vary among cell populations. Finally, transcriptome deconvolution reveals notable differences in cell composition between the two cohorts.

The convergence of signalling pathways between the two cohorts, despite the small overlap between the DEGs in both cohorts, shows that aspirin modulates robustly conserved biological processes. Aspirin is known to act on many mechanisms identified in our study such as inflammation, lipid metabolism, platelet signalling, cell proliferation, and hypoxia ^35,36^.

Several factors can explain the moderate overlap between the DEGs of the two cohorts, such as interindividual biological variability, including differences in age, comorbidities, dose or duration of aspirin exposure, that could influence the transcriptional response. However, clearly the major key element in interpreting the observed differences at the gene level is clearly provided by the deconvolution analysis. We observe important differences in the composition of the samples from both cohorts. First cohort is enriched in SCT (58%), while second cohort has a small proportion of SCT (9.3%) and is enriched in VCT (19%) and EVT (20%). Thus, if certain cell populations are over-or under-represented in a cohort, the genes specific to or enriched within them may appear differentially expressed, even in the absence of intrinsic modulation. This observation underscores the importance of considering cell composition when analysing bulk microarray or RNA-seq data. The fact that the pathways are similar across cohorts, despite these differences, suggests nevertheless that the effect of aspirin is global and affects multiple cell types, but that the intensity of the response (or its gene targets) varies according to tissue composition.

The 116 down-regulated genes common to both cohorts likely represent the core of the transcriptional response to aspirin in the placenta. These genes could correspond to direct targets or central mediators of the anti-inflammatory and metabolic effects of aspirin. Their expression in the majority of cell types, as revealed by scRNA-Seq, reinforces the idea that they participate in a global tissue response. Two pathways appear to be strongly associated with this set of genes : **TNFα signalling via NF-kB** and **Response to Hypoxia**. Aspirin and its metabolite, salicylate,e can inhibit the IkB Kinase (IKK) and prevent NF-kB activation both in vivo and in vitro ^20,37,38^. Till date, IKK is the only COX independent target know to interact directly with aspirin or salicylic acid. NF-κB is a key transcription factor involved in many biological processes: inflammation, the immune response, cell survival, proliferation or the stress response. By inhibiting NF-kB, aspirin reduces the production of pro-inflammatory cytokines, decreases the expression of COX-2 (PTGS2, which is itself an NF-κB-dependent gene), attenuates chronic inflammatory loops, can influence pathways related to hypoxia, proliferation, and cell death (since NF-κB interacts with HIF-1α, p53, Bcl-2).

Placental hypoxia plays a central role in the pathophysiology of preeclampsia and intrauterine growth restriction, notably through the activation of HIF-1α and HIF-2α ^39^. HIF-1α expression is increased in human placentas exposed to chronic hypoxia, and its activation is associated with angiogenic and inflammatory alterations characteristic of hypertensive disorders of pregnancy ^40,41^. Similarly, HIF-2α has been identified as a negative regulator of trophoblastic syncytialization, contributing to placental dysfunction and foetal growth restriction ^42^.

Several studies indicate that aspirin interferes with the cellular response to hypoxia, particularly through modulation of the HIF-1α axis. In various pathophysiological models, aspirin has been shown to reduce HIF-1α expression or activity, thereby attenuating hypoxic transcriptional programs. In hepatocellular carcinoma, aspirin reduces HIF-1α levels leading to down-regulation of GLUT1, decreased glucose import and reduced cellular proliferation ^43^. Aspirin has also been found to decrease the expression of H19, which sequesters the HIF1α negative regulator let-7, thereby reducing HIF-1α ^44^. Aspirin supresses PDK1 expression in breast cancer stem cells in a dose dependent manner via the HIF-1α signalling ^45^. Currently there are no studies showing that aspirin can directly alter HIF-1α expression or function in the human placenta. However, multiple studies have shown that NF-kB is a critical transcriptional activator of HIF-1α and NF-κB is required for HIF-1α accumulation under hypoxia ^46,47^. Loss of IKK-β (the kinase activating NF-κB) prevents proper HIF-1α induction and reduces expression of HIF-1α target genes such as VEGFA. Finally, NF-κB and HIF-1α are tightly interconnected in hypoxic and inflammatory environments, sharing regulators such as prolyl hydroxylases (PHDs) ^48^. Thus, Aspirin through its inhibitory effect on NF-kB could regulate hypoxia. Furthermore, aspirin influences metabolic processes closely linked to hypoxia, notably through AMPK activation and modulation of energy substrate utilization, which may interfere with the metabolic adaptation orchestrated by HIF-1α. Taken together, these data suggest that aspirin exerts a regulatory effect on the hypoxic response, which may contribute to explain the transcriptomic signatures observed in our study. Using the Rummagene tool of EnrichR we discover that the 116 gene set of downregulated genes under aspirin treatment overlaps highly significantly (adjusted p value 5.12 10^-17^, silhouette score = 0.43) with the dataset GSE236475, from ^49^, that identifies adrenomedullin as a possible therapeutic tool to overcome early onset preeclampsia, and this in the context of the core of trophoblast differentiation which is the fusion generating the syncytiotrophoblast ^50^. Another highly significant score (adjusted p value 1.04 10^-16^, silhouette score = 0.037) is found with GSE158901 ^51^ an article that studies the transition from human trophoblast stem cells to syncytiotrophoblast. These two studies indicate that besides the anti-inflammatory effect of aspirin, this molecule plays an active role in the mechanism of trophoblast differentiation. The gene list identified could then at least partly account for the fact that aspirin treatment is efficient only if administered before 16 weeks of pregnancy. Another strong correlation (p= 3.05 10^-62^) is found with data from ^52^ that posits the function of the Hippo pathway in differentiation, a cascade crucial for trophoblast differentiation ^53^. Again, this marks the connection, beyond the anti-inflammatory effects, between aspirin and trophoblast differentiation.

Amongst the gene where the aspirin effect is different when the cells fuse, it is interesting to notice that LRRC1 has not been previously formally analyzed in the context of preeclampsia; it is known are well expressed in the trophoblast, especially in the VCT, but also in the SCT. In other contexts, it is a cell differentiation factor, connected with WNT signalling and polarity of epithelial cells. In our BeWo cells, it was down-regulated in non-fused cells, but induced in fused cells, suggesting deregulation of trophoblast lineages at all stages. CRH is well known to be important in preeclampsia ^54^ ^20^^18^, in particular in regulating NO equilibrium, pivotal in the onset of the diseas ^55–57^. Soluble Endoglin (sENG) is one of the earliest circulating markers of preeclampsia ^58^. Here is one of the first report showing the direct impact of aspirin on this protein, both at the placental level and in the trophoblast cell model, this was true in non-fused cells, essentially, with a down-regulation by aspirin, while the fused cells showed an increase of the secreted protein **(Figure 6)**.

In summary, aspirin appears to induce a partial syncytiotrophoblast differentiation program in BeWo cells and enhances selected endocrine and placental signaling pathways, with GDF15 and ENG emerging as prominent aspirin-responsive target.

There are several limits of the present study, one is the sample size that is relatively limited, another one is that we concentrate upon patients that despite early low-dose aspirin administration developed preeclampsia. Therefore, aspirin failed to prevent the development of the disease, even though it may have considerably alleviated the symptoms. One cohort was studied by RNA-seq, and the other by Clariom microarrays (Affymetrix). We nevertheless suppose that this sophisticated array cover most of the possible transcript and that the results are largely comparable ^59^.

Comparing the DEGs in both cohorts with an exhaustive list of genes modified in preeclampsia, we found that aspirin down-regulated many genes that are up-regulated in the preeclamptic placenta. However, many other genes up-regulated by preeclampsia were unaffected by the aspirin administration. The ORA of the functions of these unaffected genes show that they are involved in TNF-α signalling, Hypoxia response, Apoptosis, Epithelial-mesenchymal transition and coagulation. Therefore low-dose aspirin administration was, in these patients, unable to reverse completely the transcriptional modifications induced by preeclampsia. A better understanding of interpatient difference in the response to aspirin, also in terms of genetic variants ^60^ could be advisable in order to help defining the patients for which the treatment has the best chance of efficacy.

## Acknowledgements

IB was funded by the BioSPC doctoral school.

## Supplementary figures and tables

**Supplementary figure 1:**
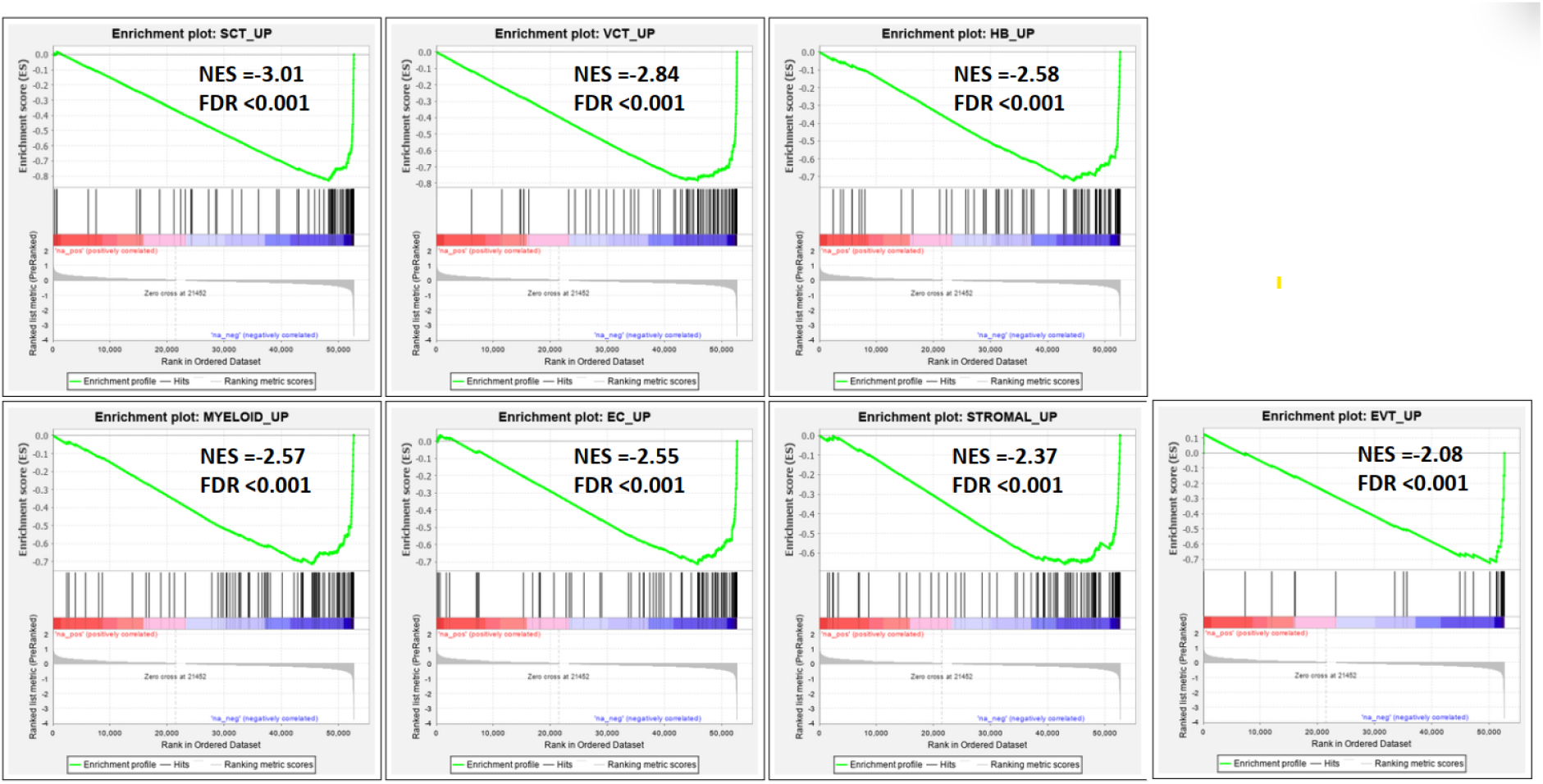
GSEA of the transcriptomic data of the first cohort against placenta cell-type specific gene sets of the preeclamptic placenta. We used a sc-RNA-Seq data set comparing preeclamptic placentas and matched non-pathological placentas at term (Admati, 2023) to extract cell-type specific signatures of deregulated genes in preeclampsia. Then GSEA was conducted against these preeclampsia cell-type specific gene sets. Very significant enrichments (FDR< 0.001) were observed for all the cell types in the negative phenotype, that is genes up-regulated in preeclampsia and down-regulated by the aspirin administration.

**Supplementary figure 2:**
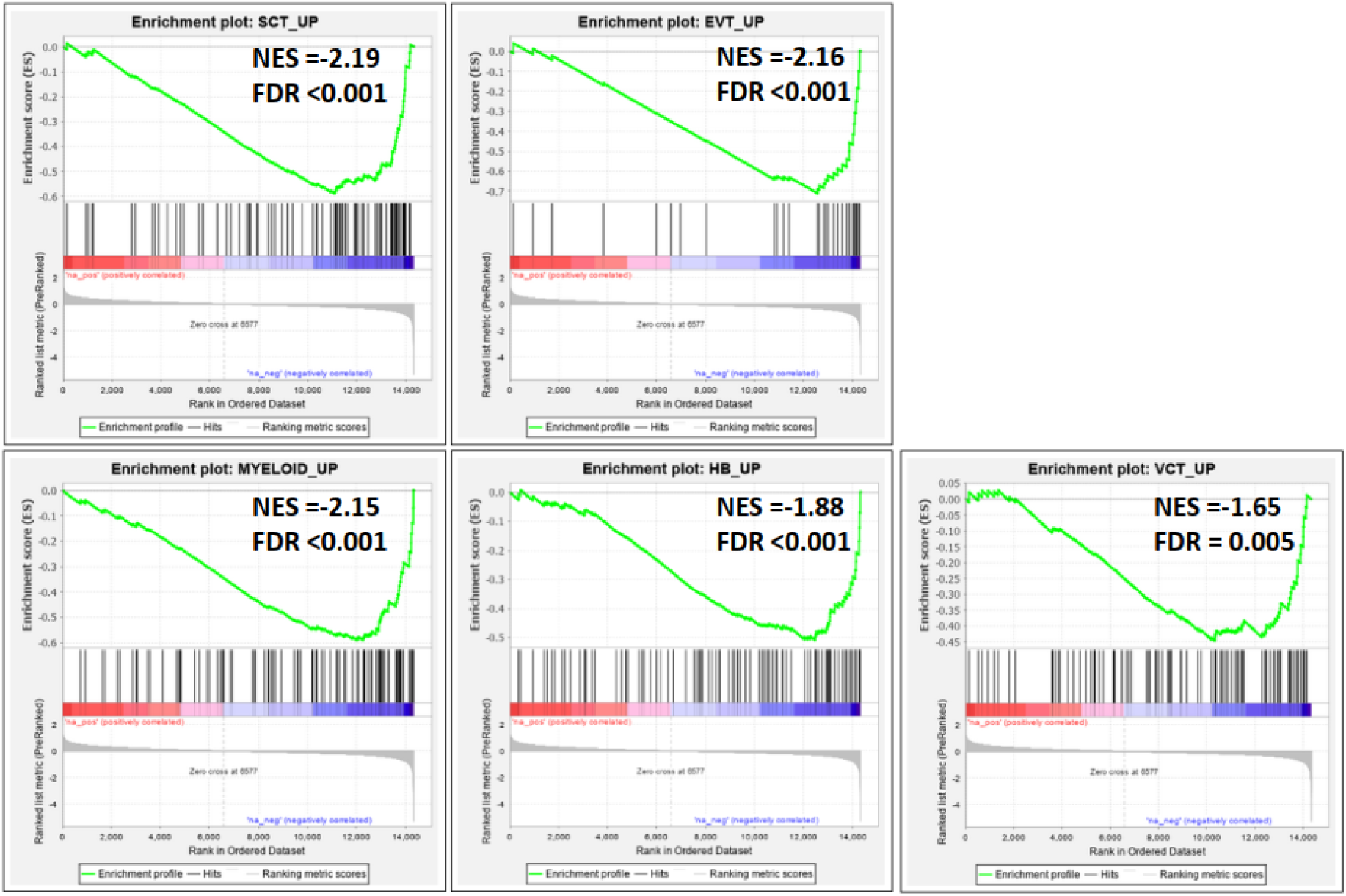
GSEA of the transcriptomic data of the second cohort against placenta cell-type specific gene sets of the preeclamptic placenta. We used a sc-RNA-Seq data set comparing preeclamptic placentas and matched non-pathological placentas at term ^23^ to extract cell-type specific signatures of deregulated genes in preeclampsia. Then GSEA was conducted against these preeclampsia cell-type specific gene sets. As for the first cohort, significant enrichments (FDR< 0.001) were observed for the cell types in the negative phenotype, that is genes up-regulated in preeclampsia and down-regulated by the aspirin administration.

**Supplementary Table 1:** Genes in the leading edge of the enriched hallmarks identified by GSEA in the first cohort.

**Supplementary Table 2**: Genes in the leading edge of the enriched hallmarks identified by GSEA in the second cohort.

**Supplementary Table 3:** The common set of down-regulated genes in both cohorts associated with enriched Hallmarks .

